# Synthetic FLS2 receptor oligomer boosts plant innate immunity

**DOI:** 10.1101/2025.01.27.634983

**Authors:** Zhiming Ma, Yi Xie, Choon-Peng Chng, Changjin Huang, Yansong Miao

## Abstract

Cell surface receptors’ gradual assembly and oligomerization are vital for controlling receptor activation and turnover. However, the spatiotemporal mechanisms of how surface receptor interactions achieve high efficiency and sustain plant immune signaling remain unclear. Here, we synthetically engineered the *Arabidopsis* pattern recognition receptor FLS2 to control its oligomerization precisely. We investigated the dynamic FLS2 nanoscale assemblies at the single-molecule level and their corresponding rewired immune signaling. Engineered FLS2 exhibits enhanced defense mechanisms in an oligomerization status-dependent manner. FLS2 dimerization significantly enhances immune responses, while the over-assembled tetrameric version impairs receptor endocytosis, disrupting its timely turnover and weakening sustained immune signaling. Our results reveal precise control of immune receptor assemblies for initial activation and long-lasting immune signaling, offering insights for engineering plant defense receptors.

## Introduction

Plants rely on innate immunity to confer resistance against pathogen attacks. Perception of pathogen- or microbe-associated molecular patterns (P/MAMPs) through the plasma membrane (PM) localized pattern recognition receptors (PRRs) initiates pattern-trigged immunity (PTI) (*1, 2*). Upon recognizing pathogenic signal, plant PRRs form complexes immediately with their co-receptors and activate downstream immune cascades by directly phosphorylating their associating partners such as receptor-like cytoplasmic kinases BOTRYTIS-INDUCED KINASE 1 (BIK1) for triggering reactive oxygen species (ROS) burst, or activate the mitogen-activated protein kinase (MAPK) cascade (*3-5*). However, the spatial and temporal assembly of resting-state immune regulators, their stepwise activation to ensure rapid and robust responses, and their efficient turnover for sustained immune perception remain poorly understood.

Upon perception of pathogenic signal, a functional assembly of receptors, co-receptors, and their associated signal molecules is critical for initiating immune signaling and systematic coordination of effective defense mechanisms. Over the past few years, our understanding of functional receptor assembly in plants has been significantly advanced by studying intracellular nucleotide-binding and leucine-rich repeat receptors (NLRs). Plant NLRs commonly oligomerize and assemble multi-protein complexes with fixed stoichiometries, termed resistosome, in response to pathogen effector proteins. Those resistosomes directly function as molecular hubs to initiate the downstream immune responses or act as holoenzymes to produce the secondary messages that enable signal amplification (*6, 7*). Oligomerization thus serves as a key regulatory step, transforming molecular recognition events into robust molecular or physiological outputs. Plant PRRs are leucine-rich repeat (LRR)-receptor kinases (RKs) or LRR-receptor proteins (RPs) featured by possessing a cytosolic kinase domain or constitutively binding with another LRR-RKs (*8*), respectively. The kinase domains of those LRR-RKs can directly mediate trans-phosphorylation with their co-receptors and further phosphorylate the downstream signaling molecules to initiate immune signaling (*3, 9*). Interestingly, oligomerization has also been recognized as a key mechanism in kinase signaling. It has been reported that phase separation or clustering of kinases into condensates can amplify or fine-tune kinase activity by concentrating enzymes and substrates, facilitating efficient signal transduction (*10, 11*). This spatial organization allows for rapid signal propagation and robust responses to stimuli both *in vivo* and *in vitro (10, 12)*. Indeed, PRR oligomerization upon activation has also been observed in plant cells. The FLAGELLIN SENSING 2 (FLS2), a well-studied plant PRR, has been shown to exhibit increased homo-association within minutes after its ligand-flg22 elicitation, as demonstrated through coimmunoprecipitation and image-based approaches (*13, 14*). While it is well-known that flg22 triggers robust heterodimerization of FLS2 and BRI1-ASSOCIATED KINASE 1 (BAK1) (*15*), the heterodimeric FLS2/BAK1 complexes could associate into larger complexes with varied stoichiometry, likely comprising two FLS2 and two BAK1 molecules (*16*). However, the regulatory mechanisms behind different types of plant PRR oligomerization and their impact on PTI signaling remain unclear.

Here, we synthetically engineered resting-state FLS2 in *Arabidopsis* to control its oligomeric state precisely. Using quantitative single-molecule analysis in living cells and monitoring immune signal transduction, we identified dimeric FLS2 as the optimal oligomeric form. This configuration enhances defense against bacterial infection by boosting receptor activation while maintaining robust endocytosis-mediated turnover, ensuring sustained innate immune responses.

## Results

### Association of resting-state FLS2 receptors upon ligand elicitation on the PM

We first applied high temporal-resolution imaging to characterize the nature of FLS2-FLS2 association in real-time following ligand elicitation in *Arabidopsis*. To avoid any weak dimerization from fluorescent protein, we generated *pFLS2∷FLS2-3×myc-msfGFP* (FLS2-mG) and *pFLS2∷FLS2-3×myc-mCherry* (FLS2-mCh), which were transiently co-expressed in *N. benthamiana* leaves (Fig. 1A). Under fluorescence lifetime imaging microscopy (FLIM), we observed an apparent decrease in the lifetime of FLS2-mG within 5 min after flg22 application (Fig. 1, B and C), which was not attributed to the homo-FRET in between FLS2-mG molecules (*14*), as such interactions are undetectable by FLIM imaging (Fig. S1, A and B). These results suggest an increased association between individual resting-state FLS2 receptors upon ligand elicitation.

**Fig. 1.**
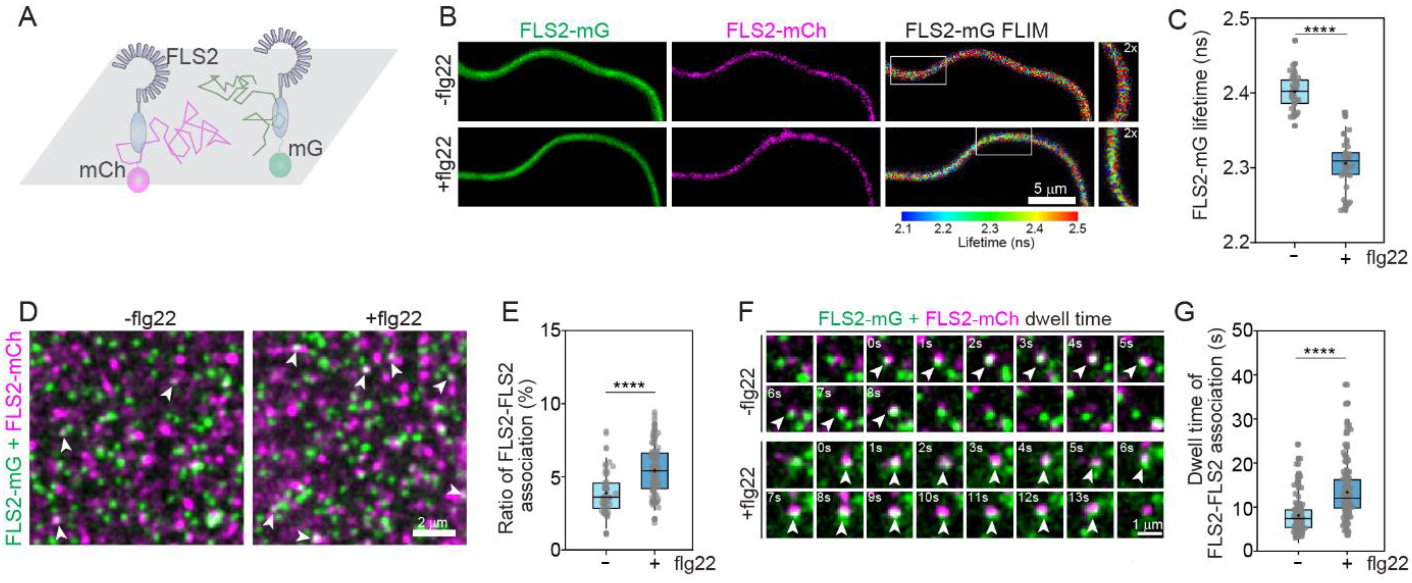
FLS2 receptor clustering upon elicitation. (**A**) Schematic illustration of dual-colour tagging and imaging for FLS2 receptor. *pFLS2-FLS2-3×Myc-msfGFP (mG)* and *-mCherry (mCh)* were transiently expressed in *N. benthamiana* leaves. Signals were recorded by either FLIM imaging or VA-TIRFM for single particle dynamic imaging. (**B**) FLIM images for FLS2-mG (donor) in the presence of FLS2-mCh (acceptor) with or without elicitation by 20 nM flg22 for 5 min. Intensity images of FLS2-mG/-mCh and the lifetime image for FLS2-mG are shown. The colour bar indicates the corresponding lifetime. Enlarged images show 2 x zoomed view of the white boxes. Scale bar, 5 μm. (**C**) Quantification of FLS2-mG lifetime in (B), n = 40 and 37 ROIs from left to right. (**D**) Overlaid images of FLS2-mG/mCh single particles with or without elicitation by 20 nM flg22 in 5 min. FLS2-mCh images were processed using a deconvolution algorithm (see Fig. S1C). Arrowheads mark FLS2-FLS2 association. Scale bar, 2 μm. (**E**) Quantification of the ratio of FLS2-FLS2 association in (D), the number of particles showing association was normalized by the total number of FLS2 single particles. n = 36 ROIs (area of 100 μm^2^). (**F**) Real-time recording of FLS2 single particle association and disassociation. Arrowheads mark two particle associations with indicated dwell time. Scale bar, 1 μm. (**G**) Quantification of FLS2-FLS2 association dwell time in (F). n = 117 and 123 of associations from left to right. Box plots in (C), (E) and (G) display the mean and median (cross symbols and center bars), upper and lower quartiles, and whiskers representing SD. Significant differences were determined by unpaired *student’s* test (****p ≤ 0.0001).

To obtain detailed insight into FLS2-FLS2 association at a single receptor level on the PM, we next monitored the dynamic diffusion of FLS2-mG/mCh single particles (probably single molecules) under variable angle-total internal reflection fluorescence microscope (VA-TIRFM). We observed a few FLS2-mG/mCh particles showing colocalization at their resting states (Fig. 1D, Fig S1C). Such association has a significantly higher probability than incidental overlapping of random particles at similar densities on a two-dimensional surface in a simulation system (Fig. 1E; Fig. S1, D and E), where the two-particle association rate scales with the relative inter-particle attraction. Notably, the proportion of FLS2-mG/mCh particles showing colocalization was increased after flg22 elicitation (Fig. 1, D and E; Fig. S1C), which is in line with the FLIM image data (Fig. 1, B and C) and bulk coimmunoprecipitation showing increased FLS2-FLS2 association after ligand elicitation (*13*). Real-time imaging revealed that FLS2-mG/mCh clusters, once formed, were transient and disassembled within seconds (Fig. 1F, Movie S1). The dwell time of these FLS2-mG/mCh clusters was significantly increased following flg22 elicitation (Fig. 1, F and G; Movie S2). Collectively, those data clearly demonstrate ligand elicitation triggers increased FLS2-FLS2 association. Therefore, we hypothesize that prolonged, dynamic FLS2-FLS2 associations act as activation factors, promoting the formation of functional FLS2 assemblies that boost innate immune activation.

### Synthetic engineering of FLS2 for defined oligomeric states

The dynamic association and dissociation of FLS2 clusters, mixed with un-clustered resting-state FLS2, complicates the analysis of receptor clustering status and its role in immune activation. Thus, we generated transgenic *Arabidopsis* lines (in complement of *fls2*) stably expressing engineered FLS2 receptors with precise oligomerization control (Fig. 2A). Specifically, we inserted dimeric (D) and tetrameric (T) coiled-coil (CC) motifs (*17*) between FLS2 and mG in the FLS2-mG (monomeric, M) construct. Using single-particle tracking, we observed that these FLS2 variants (FLS2-M/D/T-mG) exhibited distinct intensity and diffusion dynamics on the PM at their resting states (Fig. S2, A and B), suggesting potential differences in their oligomerization.

**Fig. 2.**
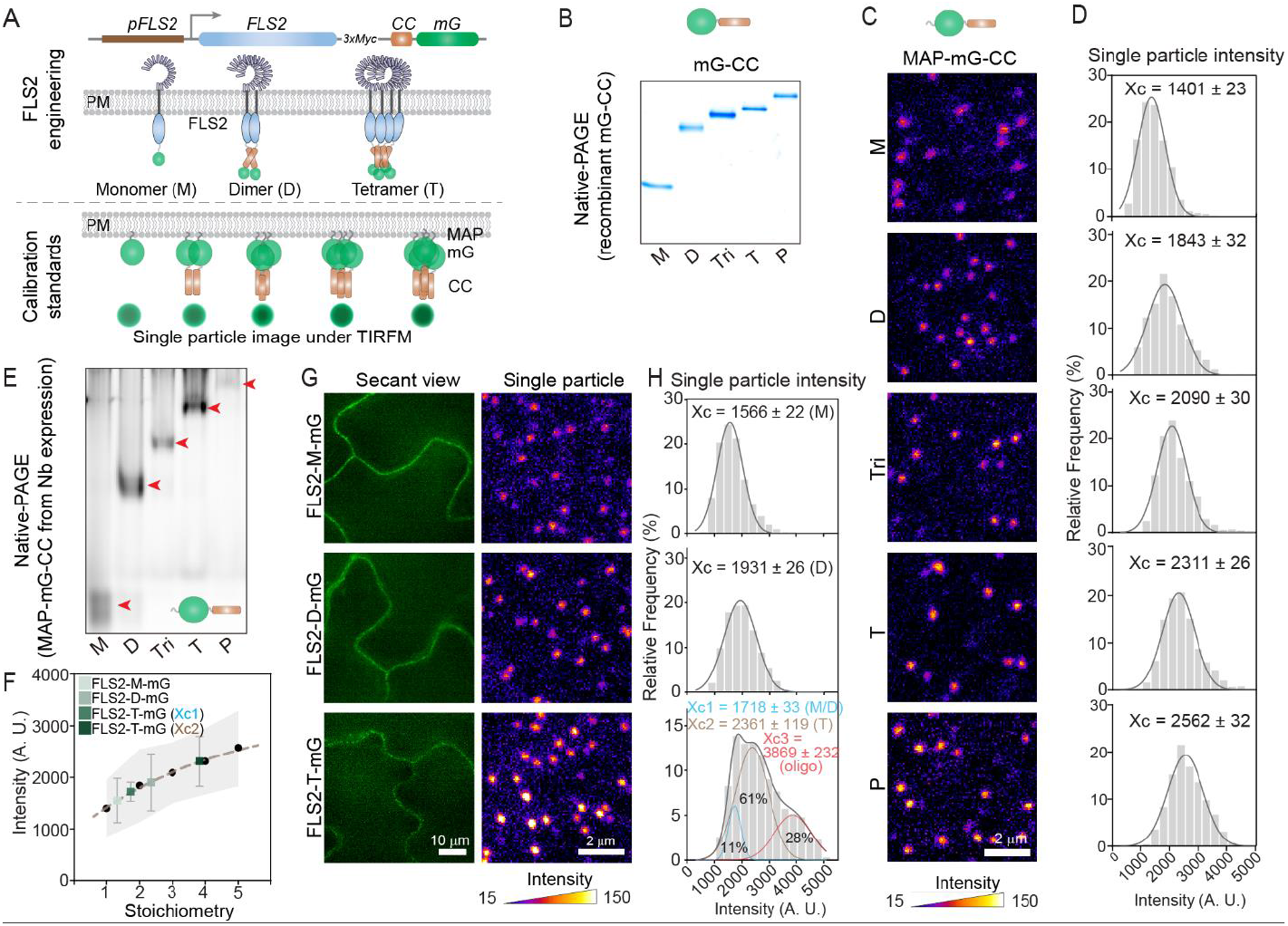
FLS2 engineering for a defined FLS2 clustering. (**A**) Schematic illustration of FLS2 engineering to reach a precisely controlled FLS clustering. FLS2 was fused with a Coiled-coil (CC) domain followed by mG. CC domains assemble to multimers: Dimer (D) or Tetramer (T). Construct without CC domain was designated as Monomer (M). Single particle calibration standards were designed as CC domains (forming dimer to pentamer) fused with mG and an N-terminal MAP peptide for membrane targeting. (**B**) Coomassie blue-stained SDS-PAGE gel image of recombinant mG-CC proteins with CC valencies: Dimer (D), Trimer (Tri), Tetramer (T) or Pentamer (P). mG without a CC domain was designated as Monomer (M). (**C**) Single particle images of MAP-mG-CC calibration standards transiently expressed in *N. benthamiana* leaves, as in (A). The colour bar indicates the intensity. Scale bar, 2 μm. (**D**) Plots of single-particle intensities for MAP-mG-CCs in (C), fitted to Gaussian distributions. Xc represents centre peak values used as calibration standards for determining FLS2 oligomeric status *in vivo*. n=1537/1535/1483/1376/1528 for MAP-mG-M/D/Tri/T/P, respectively. (**E**) Native-PAGE gel image for MAP-mG-CCs from *in vivo*. Total proteins from *N. benthamiana* leaves were extracted and separated. Native GFP signals were captured under UV. Red arrowheads mark corresponding MAP-mG-CC protein bands. (**F**) Plot of mG single-particle intensities as a function of mG stoichiometries determined by valencies in (C). Black dots or coloured squares indicate the intensity peaks while shaded area or error bars represent full width at half maximum from Gaussian fits in (C) or (H). A single exponential function fitting of intensity peaks from (C) generated the calibration standard curve (dash line), which was aligned with those of FLS2-CC-mGs to determine FLS2 clusters’ stoichiometries. (**G**) Secant view and single-particle images from cotyledon epidermal cells of 5-day-old *Arabidopsis* stable transgenic FLS2-CC-mG seedlings. The colour bar indicates intensity. Scale bar: 10 μm (left) and 2 μm (right). (**H**) Quantification of FLS2-CC-mG single-particle intensities in (G), subjected to Gaussian distribution fitting with single or multiple peaks. n=1508/1642/1672 for FLS2-M/D/T-mG, respectively. Xc represents centre peak values, which correspond to FLS2-CC-mG oligomeric status indicated in brackets aligned within (F).

To precisely characterize the oligomeric states of engineered FLS2 receptors, we developed a single-particle intensity-based calibration curve to measure the oligomerization of any fluorescent fusion protein on the PM. This system uses monomeric to pentameric mG-CC tags combined with single-particle imaging (*17*) (Fig. 2A), whose precision in particle intensity analysis is independent of protein-expression level and particle density on the PM. Firstly, all mG-CC standards could be assembled in correct states *in vitro* as reported (*17*), which were characterized using recombinant proteins (Fig. 2B and Fig. S2C). Next, all these mG-CCs were targeted to the PM by fusing with a peptide with myristylation and palmitoylation (MAP) (*18*) (Fig. 2A). Through transient expression in *N. benthamiana* and single-particle imaging via VA-TIRFM, we demonstrated that those cell surface mG variants exhibited progressively increasing single-particle intensities corresponding to their valencies, from monomer to pentamer (Fig. 2, C and D). This was further validated by native PAGE gel separation of the extracted mG variants from *N. benthamiana* cells (Fig. 2E). Next, we set the calibration standard by quantifying the single-particle intensities of the MAP-CC-mGs (Fig. 2, D and F). By applying FLS2-M/D/T-mG intensity onto the MAP-mG-CC standard curve (Fig. 2, F-H), we measured the stoichiometries of all engineered FLS2 clusters. As a result, FLS2-M-mG and FLS2-D-mG predominantly assemble as monomer and dimer, although their intensity peaks were slightly higher than those of standard monomeric- and dimeric-mG on the PM (Fig. 2, F-H), respectively. Such discrepancy could be attributed to a small fraction of FLS2 also undergoing self-association at resting-state (Fig.1, D and E). In contrast, FLS2-T-mG displayed a mixed population, with the majority (61%) forming tetramers, along with smaller fractions of lower-order oligomers (11%) and higher-order oligomers (28%) beyond the calibration range (Fig. 2, F-H). This suggests the challenge of maintaining precise oligomerization *in vivo* for highly oligomeric FLS2, likely due to its inherently weak self-association properties and unintended immobilization (Fig. S2A).

### Receptor clustering enhances PTI signaling

Next, with those engineered FLS2 receptors with defined clustering stoichiometries, we investigated a series of plant immune responses following FLS2 activation to evaluate how each engineered FLS2 influences immune activation. With a comparable FLS2 expression across FLS2-M/D/T-mG lines, we first confirmed that the FLS2 engineering did not affect plant growth (Fig. S2, D-F). Interestingly, FLS2-D-mG and FLS2-T-mG showed noticeable elevation in activating MAPK cascade and producing ROS, in which FLS2-T-mG showed relatively lower activities compared to FLS2-D-mG (Fig. 3, A-C). Consistent with the rapid PTI activation results, the growth trade-offs caused by long-lasting PTI responses indicate stronger PTI activation in FLS2-D-mG and FLS2-T-mG compared to monomeric FLS2, as evidenced by a more pronounced reduction in root growth (Fig. 3D). Overall, both FLS2-D-mG and FLS2-T-mG demonstrated greater resistance to infection by *Pseudomonas syringae* pv. tomato (*Pst*) DC3000 compared to FLS2-M-mG, with FLS2-D-mG showing the most substantial effect (Fig. 3E). Consistently, similar results were observed under infection by *Pst* DC3000 D36E (Fig. S2G), which only triggers PTI (*19*), further supporting clustered FLS2 enhances PTI responses. Our results demonstrate a higher order of FLS2 clustering, starting from the tetrameric assembly, still elevates PTI signaling but displays a less pronounced effect than dimeric-FLS2, probably because of reduced conformational flexibility at high oligomeric states.

**Fig. 3.**
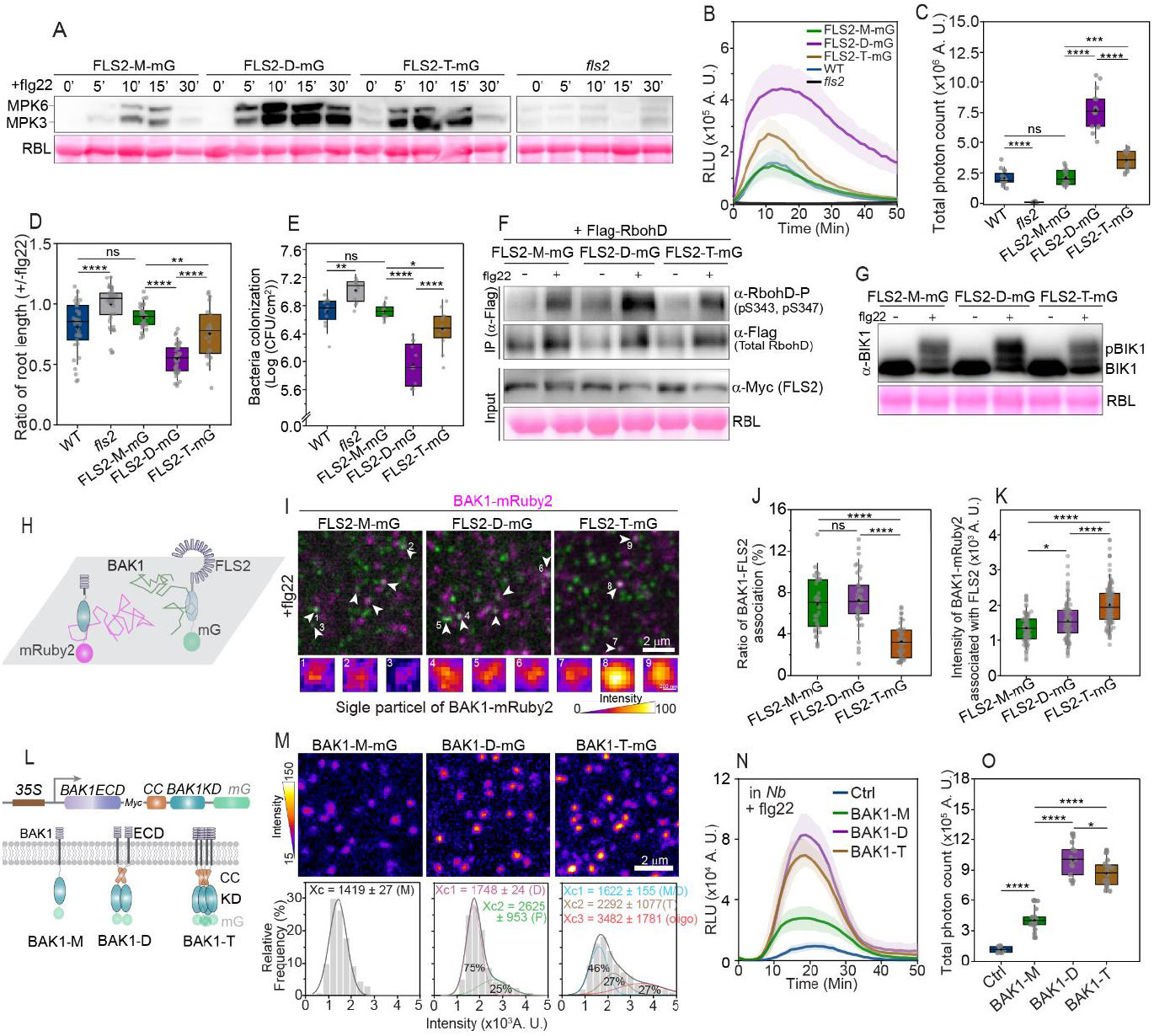
Clustering of receptor complex increases PTI signaling. (**A**) MAPK activation in *fls2* and FLS2-CC-mG *Arabidopsis* seedlings elicited by 20 nM flg22, determined over time using immunoblotting by anti-p-44/42 MAPK antibodies. Ponceau S staining of rubisco large subunit (RBL) served as a loading control. (**B**) Measurement of ROS in leaf discs from 5-week-old *Arabidopsis* lines elicited by 20 nM flg22. Relative luminescence units (RLUs) were monitored in real-time. Solid lines and shaded area represent mean ± SD from multiple leaf discs. (**C**) Quantification of total photon readings (5-30 min) in (B), representing the total ROS generation in indicated genotypes. n = 16, 16, 15, 16, 13 of leaf discs from left to right. (**D**) Quantification of root length with or without flg22 treatment. WT, *fls2* and FLS2-CC-mG *Arabidopsis* seeds were grown vertically on 1/2 MS medium supplemented with or without 1 μM flg22 for 5 days before measurement. n= 35, 42, 36, 43 and 23 of individual seedlings from left to right. See also Fig. S2, D-F. (**E**) Quantification of internal bacteria colonization at 3 days post-infection. 1 × 10^6^ CFU·mL^−1^ of *Pst* DC 3000 was injected into the rosette leaves of 4-weeks-old WT, *fls2* and FLS2-M/D/T-mG *Arabidopsis* seedlings. n=12 replicates per genotype. Error bars = SD. (**F**) RbohD phosphorylation in the presence of FLS2-M/D/T-mG co-expressed with RbohD-flag in *N. benthamiana* leaves and elicited with or without 20 nM flg22 for 10 min. RbohD was enriched using anti-flag beads and analyzed by phosphorylation-specific antibody for S343/S347. Ponceau S staining of RBL served as a loading control. (**G**) BIK1 phosphorylation, as indicated by slower migrated bands, was detected with anti-BIK1 antibodies. 10-day-old FLS2-M/D/T-mG *Arabidopsis* seedlings were treated with 20 nM flg22 for 20 min before sample collection and protein separation. Ponceau S staining of RBL served as a loading control. (**H**) Schematic of dual-colour tagging and imaging for FLS2-CC-mGs and BAK1-mRuby2 transiently expressed in *N. benthamiana* leaves and imaged using VA-TIRFM. (**I**) Representative overlap images of FLS2-M/D/T-mG and BAK1-mRuby2 single particles captured simultaneously as in (H) after 20 nM flg22 elicitation in 5 min. White arrowheads indicate FLS2-M/D/T-mG and BAK1-mRuby2 association. Extracted single particles of BAK1-mRuby2 from numbered marks were displayed in lower panels. Colour bar indicates intensity. Scale bars, 2 μm and 200 nm. (**J**) Quantification of FLS2-BAK1 association ratios in (I). The number of associations was normalized to the total FLS2 and BAK1 particles. n = 36, 42 and 42 of ROIs from left to right. (**K**) Quantifying the single-particle intensities for those BAK1-mRuby2 associated with FLS2-M/D/T-mG after flg22 elicitation. n = 80,113 and 123 of single particles from left to right. (**L**) Schematic of BAK1 engineering for clustering. The CC domains (D and T) were integrated in between the transmembrane domain and the kinase domain (KD) of BAK1. The extracellular domain (ECD) is labelled. Construct without CC domain was designated as M. C-terminal mG fusion was used for imaging but omitted in ROS assays. (**M**) Single-particle imaging of BAK1-M/D/T-mG on the PM of *N. benthamiana* leaves by transient expression. The single-particle intensities were quantified and fitted to single- or multi-peak Gaussian distributions. Xc values were aligned with calibration curves (Fig. 2F) to determine BAK1 cluster stoichiometries, as shown in brackets. n = 474, 495 and 923 from left to right. Scale bars, 2 μm. (**N**) Measurement of ROS in *N. benthamiana* leaf discs with comparable expression of BAK1-M/D/T or empty vector (Ctrl) after 20 nM flg22 elicitation. RLU was monitored in real time. Solid lines and shaded area represent mean ± SD. (**O**) Quantification of total photon reading (5-30 min) in (N), representing total ROS generation under BAK1 expression. n = 8, 16, 14 and 16 leaf discs from left to right. Box plots in (C), (D), (J), (K) and (O) indicate mean (cross symbols) and median (centre bars), quartiles (box limits), and SD (whiskers). Significant differences were determined via one-way ANOVA with multiple comparisons (****p ≤ 0.0001, ***p ≤ 0.001, **p ≤ 0.01, *p ≤ 0.05, ns = not significant).

Given many plant LRR-RKs share similar structure conformation and downstream signaling components (*3*), we were motivated to ask whether the above phenotypes on FLS2 clustering can also be observed on other typical surface immune receptors. We then engineered another LRR-RK, *Arabidopsis* PEPR1 precepting damage-associated molecular patterns (*20*), through the same strategies applied on FLS2 (Fig. 2A). Interesting, using *N. benthamiana* transient expression system (Fig. S2, H and I), the ligand pep1 triggered significantly higher ROS production in the presence of clustered PEPR1 receptors, in which PEPR1-D-mG also showed more prominent effect than PEPR1-T-mG (Fig. S2, J and K). These data suggest receptor clustering, particularly the dimeric version, greatly enhances plant PTI signaling.

We next sought to understand the underlying mechanism by investigating the receptor activation-mediated phosphorylation and activation of NADPH oxidase RESPIRATORY BURST OXIDASE HOMOLOGUE D (RbohD) on the PM for ROS production (*21, 22*). We first examined the flg22-mediated RbohD phosphorylation at S343/S347 (*23*) in different synthetically engineered FLS2 receptors in *N. benthamiana* leaves. At a comparable level of expression in *N. benthamiana*, FLS2-D-mG and FLS2-T-mG exhibited elevation in ROS production (Fig. S3, A-C) in response to flg22, similar to in *Arabidopsis* (Fig. 2, B and C). In addition, flg22 triggered significantly higher phosphorylation of RbohD (S343/S347) in FLS2-D-mG than FLS2-M-mG (Fig. 3F), but not that obvious in FLS2-T-mG (Fig. 3F), although FLS2-T-mG still generated relatively higher ROS than FLS2-M-mG (Fig. S3, A-C). We speculated that the changes in RbohD phosphorylation in FLS2-T-mG might be modest and thus undetectable by immunoblot or could involve additional phosphorylation sites beyond S343/S347(*21, 24*). Consistently, similar results on RbohD (S343/S347) phosphorylation were also observed with the set of engineered PEPR1 oligomers, particularly in PEPR1-D-mG (Fig. S3D).

RbohD S343/S347 are directly phosphorylated by the receptor-like cytoplasmic kinase BIK1 upon flg22-triggered phosphorylation and activation of BIK1 (*21, 25-27*). Given that clustered FLS2 might induce a local accumulation of BIK1 as BIK1 directly associates with FLS2 (*26*), we tested if engineered clustering of BIK1 will also have a similar effect in regulating ROS production like FLS2 clustering. However, we did not observe a clear difference in flg22-triggered ROS production across different BIK1 oligomeric states (Fig. S3, C and E-G), suggesting oligomeric BIK1 does not change BIK1 activity on activating RbohD. We next also compared the BIK1 phosphorylation level in FLS2-M/D/T-mG *Arabidopsis* lines upon flg22 stimulation. Interestingly, the slower migrated phosphorylated bands of BIK1 were prominently increased in FLS2-D-mG compared to FLS2-M-mG, although not in FLS2-T-mG (Fig. 3G). In summary, our results show that the engineered FLS2-D-mG significantly enhances the PTI signaling cascades shown by elevated BIK1 and RbohD phosphorylation, a level of enhancement that cannot be achieved by engineering BIK1 alone.

BIK1 is directly phosphorylated by BAK1, whose activation depends on the ligand-triggered FLS2-BAK1 complex formation (*26, 27*). We asked if engineered-FLS2 changes its ability in binding with BAK1 in *Arabidopsis*. Notably, FLS2-M-mG and FLS2-D-mG showeda similar binding affinity with BAK1, as indicated by Co-IP experiments (Fig. S3H). In contrast, FLS2-T-mG exhibited a significantly diminished interaction with BAK1 (Fig. S3H). Nevertheless, FLS2-T-mG *Arabidopsis* retained comparable BIK1 phosphorylation levels and slightly higher ROS production in response to flg22 elicitation than FLS2-M-mG (Fig. 3, B, C and G). This indicates that FLS2-T-mG has a mixed effect on different steps of the signaling cascade, in contrast to the consistent enhancement observed at every step with FLS2-D-mG. In addition, we also applied a single-particle imaging approach to study the flg22-triggered BAK1 association with each engineered oligomeric FLS2 receptor (Fig. 3H). We found that FLS2-M/D-mG showed a similar tendency to associate with BAK1-mRuby2, while FLS2-T-mG displayed a noticeably lower tendency (Fig. 3, I and J), consistent with Co-IP results (Fig. S3H). Interestingly, the signal intensity of BAK1-mRuby2 associated with FLS2-CC-mGs within the FLS2-BAK1 complex increased depending on the oligomeric status of FLS2 (Fig. 3, I and K). This suggests enhanced recruitment of BAK1 into individual FLS2 clusters when FLS2 forms a higher oligomer. We thus speculated that the elevated ROS production in FLS2-T-mG compared to FLS2-M-mG could be attributed to those clustered BAK1, though at low abundancy (Fig. 3, I-K). To test this hypothesis, we next engineered BAK1 by integrating the CC motifs in between its transmembrane domain and kinase domain (BAK1-M/D/T) (Fig. 3L) without other tags at the C-terminus, because tagging BAK1 at its C-terminal partially abolishes BAK1 function in PTI signaling although its interaction with FLS2 can be kept (*28*). In addition, when we determined the stoichiometries of BAK1 clusters, a mG was further added at the C-terminus (BAK1-M/D/T-mG) (Fig. 3L), which allowed us to do single-particle image and alignment with the calibration curve. We found BAK1-M-mG was clearly assembled to monomer, and BAK1-D-mG displayed dominantly (75%) as a dimer and partially (25%) as a pentamer. Whereases BAK1-T-mG again showed a multi-oligomeric assembly with around half (46%) as a monomer/dimer mixture and another half as a tetramer (27%) and higher-order of oligomer (27%). Despite the uniformed assembly in BAK1-D/T-mG, we observed flg22 application triggered higher ROS production in the presence of BAK1-D or BAK1-T clusters in contrast to monomeric BAK1-M (Fig. 3, N and O), suggesting positive functions of dimeric and tetrameric-engineered BAK1 in enhancing PTI signaling. As a co-receptor of multiple PRRs, we found engineered BAK1 also increased immune signaling triggered by DAMPs. When applying pep1 on *N. benthamiana* leaves co-expressing a similar amount of engineered BAK1-M/D/T and PEPR1-M-mG, elevated ROS productions were observed in BAK1-D/T expressing plants (Fig. S3, I-K). In summary, we showed the engineered oligomerization of resting-state PRRs and the co-receptor BAK1 significantly enhance the downstream signaling cascade upon ligand stimulation, driven by enhanced phosphorylation of intracellular signaling molecules through clustered receptor and co-receptor complexes.

### Over-assembled receptors defect in turnover and long-lasting immunity

Single-particle dynamic tracking showed that flg22 enhanced the association between resting-stated monomeric FLS2-mG and FLS2-mCh, which would not further engage more FLS2 molecules until they dissociation (Fig. 1F and Movie S2). In addition, FLS2-D-mG and FLS2-T-mG did not show increased association with FLS2-mCh upon flg22 stimulation (Fig. S4, A and B). These results suggest that the dimeric configuration is a more optimal functional form of FLS2 after its activation and therefore the engineered oligomeric FLS2 tends to bypass the need for further FLS2-FLS2 clustering. This is further evidenced by the FLIM image showing an unchanged mG lifetime in FLS2-D/T-mG co-expressing with FLS2-mCh and following flg22 elicitation (Fig. S4C). We next sought to understand the mechanisms that make the dimeric configuration more optimal than the engineered tetrameric state. One of the prominent features of clustered-FLS2 is their dynamic diffusions were highly hindered (Fig. S2A, movie S3), especially for the FLS2-T-mG, which almost were frozen on the PM (Fig.4, A and B). We therefore asked whether the dynamic turnover of FLS2 molecules after FLS2 activation is altered with clustered FLS2. Except for the constitutive endocytosis and recycling of FLS2 at its un-activated states (*29*), the ligand activated-FLS2 is ubiquitinylated and further internalized through clathrin-mediated endocytosis (*30, 31*), which was eventually transferred to the degradation pathway (*32*). Meanwhile, the de novo synthesized FLS2 is delivered back to the surface to regenerate resting-state FLS2 (Fig. 4C) (*32*). flg22 activates the endocytosis of FLS2, thereby exhibiting a shorter lifetime under VA-TIRFM imaging, indicating a faster internalization (*14, 33*) (Fig. 4, A and B). Kymograph analysis of FLS2 internalization showed that FLS2-D-mG displayed comparable efficiency as FLS2-M-mG at either resting- or flg22-simulated-state, though FLS2-D-mG exhibited relatively reduced lateral diffusion dynamics compared to FLS2-M-mG (Fig. 4, A and B; Fig. S2, A and B). In contrast, most of the FLS2-T-mG particles displayed long-lasting staying on the PM even upon flg22 stimulation, indicating retarded internalization (Fig. 4, A and B). In addition, FLS2 engagement into intracellular endosomes was robust for both FLS2-M-mG and FLS2-D-mG at 1 h post flg22 elicitation, whereas FLS2-T-mG accumulation in endosomes was only clearly observed at 2 h post flg22 elicitation and less in number (Fig. S4, D and E), suggesting the largely impaired endocytosis.

**Fig. 4.**
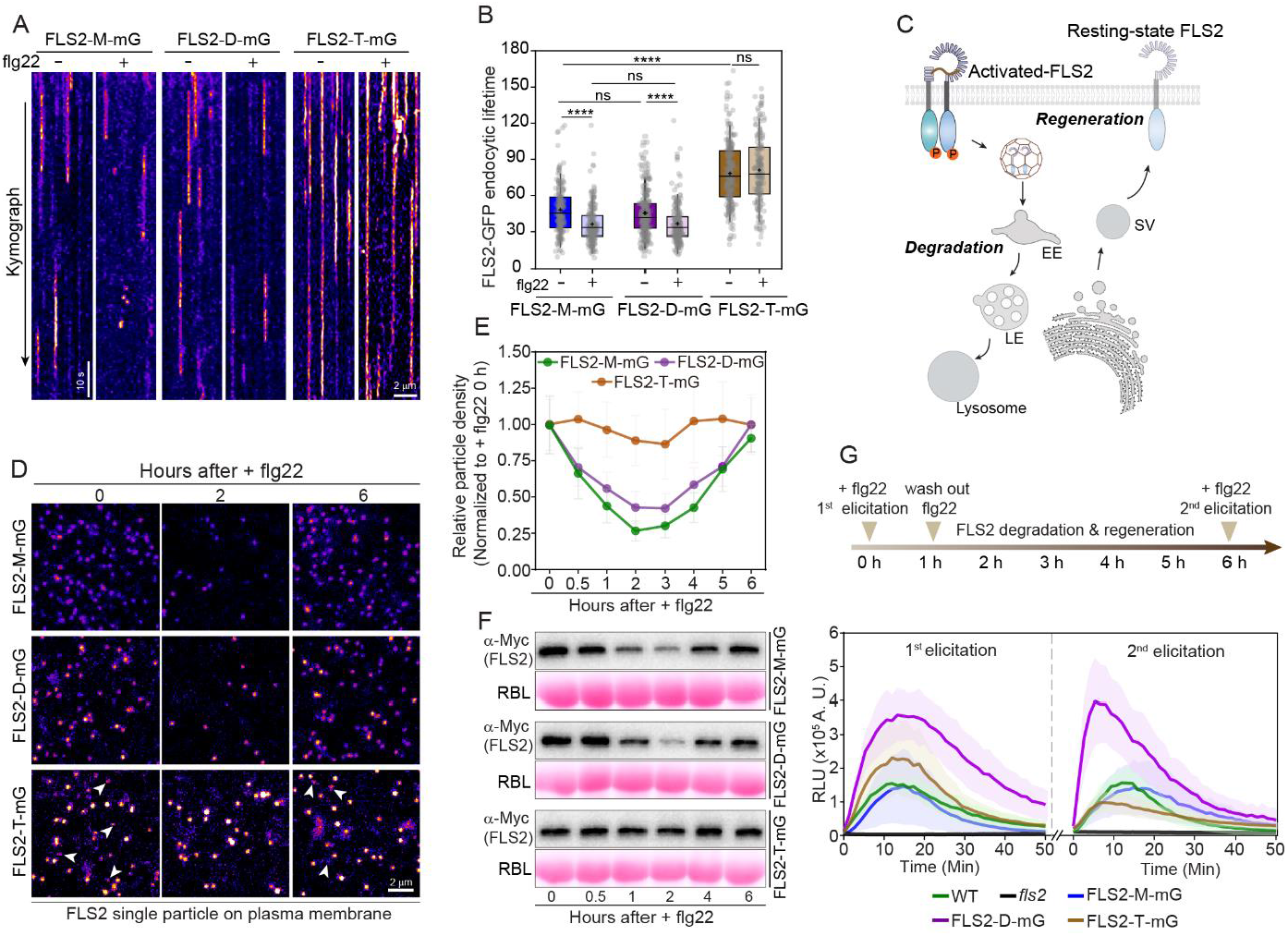
Over assembled-receptor clustering defects in long-lasting immune perception. (**A**) Kymographs showing FLS2-M/D/G-mG single-particle dynamics on the PM, recorded by the time-lapse image in the cotyledon epidermal cells of 5-day-old *Arabidopsis* seedlings with or without 20 nM flg22 elicitation for 1h. Straight lines from emerging to disappearance indicate FLS2 particles undergoing endocytic internalization. Scale bar, 2 μm. (**B**) Quantification of the FLS2 endocytic lifetime in (A). Noted that FLS2-T-mG lifetime was underestimated, as most of FLS2-T-mG particles remained immobile, as shown in (A) and Fig. S2A. Box plots indicate mean (cross symbols), median (center bars), quartiles (box limits), and SD (whiskers). n = 185, 197, 234, 227, 224 and 167 from left to right. Significant differences were determined via one-way ANOVA with multiple comparisons (****p ≤ 0.0001, ns = not significant). (**C**) Simplified model of receptor turnover after ligand elicitation. The ligand-activated receptors will be engaged into clathrin-coated vesicles and transferred to early endosome (EE), then late endosome (LE) and eventually lysosome to be degraded. Meanwhile, newly synthesized receptors in secretory vesicles (SC) are transported to the PM, regenerating resting-state receptors. (**D**) Real-time imaging of FLS2-M/D/T-mG single particles on the PM after elicitation by 20 nM flg22, as in (A). flg22 was washed out 1 h post-elicitation to avoid continuous stimulation. Arrowheads highlight low oligomeric-FLS2 of FLS2-T-mG particles that disappeared and were regenerated at 6 h. A complete image set with more time points is shown in Fig. S4F. Scale bar, 2 μm. (**E**) Quantification of single-particle density (normalized to 0 h post elicitation) in (D) and Fig. S4F. n ≥ 35 of ROIs in each time point. Error bar = SD. (**F**) Immunoblot analysis of bulky FLS2 protein level (detected by ani-Myc) at various time points post-flg22 elicitation as in (E). Leaf discs from 5-week-old FLS2-M/D/G-mG *Arabidopsis* seedlings were treated with 20 nM flg22, which was washed out at 1 h. Ponceau S staining of rubisco large subunit (RBL) was used as a loading control. (**G**) Two-step ROS measurement in the leaf discs from 5-week-old *Arabidopsis* lines elicited with 20 nM flg22. The ligand was washed out at 1 h post-first elicitation. A second elicitation occurred at 6 h post-first elicitation when receptors are regenerated, as shown in (D-F). Relative luminescence units (RLU) were monitored in real-time. Solid lines and shaded area represent mean ± SD. n = 6, 8,13,12 and 9 of leaf discs for WT, *fls2*, and FLS2-M/D/G-mG, respectively.

Next, we aimed to investigate the roles of spatiotemporally regulated receptor endocytosis and regeneration in immunity. We first followed receptor dynamic over time upon flg22 elicitation. FLS2-M/D-mG particles, but not most of FLS2-T-mG, disappeared at 2 h after flg22 application and then progressively recovered eventually to a comparable level as before flg22 application after 6 h post adding flg22, demonstrating dynamic turnover of FLS2 from internalizing activated receptors to replenish resting-states ones (Fig. 4, D and E; Fig. S4F). Consistently, immunoblot analysis revealed similar oscillatory homeostasis in the protein levels of FLS2-M-mG and FLS2-D-mG following flg22 elicitation. At the same time, no such detectable changes were observed in FLS2-T-mG (Fig. 4F). These results suggest tetrameric- and higher order of FLS2 clusters failed to be degraded. Therefore, there is no regeneration of non-activated FLS2. Then, we hypothesized that the dynamic turnover of FLS2-M-mG and FLS2-D-mG is crucial for sustaining continuous elicitation, thereby supporting long-lasting PTI responses. We thus measured the ROS production by doing a two-step flg22 elicitation in FLS2-M/D/T-mG *Arabidopsis* to evaluate the consequence of the hindered FLS2 degradation in FLS2-T-mG (Fig. 4G). One hour after the first elicitation, flg22 was washed off to prevent continuous stimulation. The second elicitation was performed six hours after the first. Interestingly, both FLS2-M-mG and FLS2-D-mG, but not FLS2-T-mG, generated a similar level of ROS upon the second elicitation, as the first elicitation, utilizing regenerated resting-state FLS2 molecules (Fig. 4G). In contrast, ROS production was significantly reduced during the second elicitation in FLS2-T-mG (Fig. 4G), which is reminiscent of earlier studies showing that activated FLS2 cannot be re-elicited for immune signaling despite it remains undegraded (*32*). In conclusion, the dynamic turnover of activated-FLS2 is essential for maintaining long-lasting PTI. Over-assembled FLS2, such as FLS2-T-mG, impairs its internalization and degradation, thereby compromising the sustained immune perception.

## Discussion

Plant PRR clustering during immune activation has been reported in earlier studies (*13, 14, 16, 34*). However, the precise stoichiometric controls of plant PRRs per se or in combination with their co-receptors and the resulting functional outcomes are not clear. While structural studies have well documented that ligand binding induces heterodimerization of receptor and co-receptor, the *in vivo* resting state of receptors and the precise mechanisms by which they transition into spatiotemporally regulated complexes to coordinate dynamic behaviors after ligand perception remain enigmatic. Such precise characterization has been extensively investigated in mammals. The Toll-like receptors (TLRs), sharing structural and functional similarities with plant PRRs (*9*), undergo dimerization upon activation to induce conformational changes that are essential for the initiation of immune signaling (*35*). TCR activation also induces high-order assembly of signaling components by molecular condensation to trigger downstream signaling (*36*). Human formyl peptide receptor, a typical G protein-coupled receptor (GPCR), has a dynamic monomer-dimer equilibrium in living cells either before or after ligand binding (*37*), of which the ligand-specific dimeric conformation is vital for downstream functional responses (*38*). The epidermal growth factor receptors (EGFRs) convert from a monomeric-to dimeric-form upon binding ligands to release the dimerization arms and thus activate their kinase domains (*39*). However, in plant cells, it remains unclear which assembly states of surface receptors optimize resting state activity for maximal signal transduction, and which oligomeric forms most effectively manage the entire signaling process from initial ligand-induced activation to the restoration of receptor function through dynamic turnover.

Here, through quantitative single-particle image analysis, we demonstrated that FLS2 is present as a monomer in a resting state on the PM before activation. Our single receptor investigation is consistent with previously reported bulk assays demonstrating both FLS2 and BAK1 form neither homomers nor heteromers before flg22 elicitation (*16*). Upon application of flg22, in a time window of tens of seconds, we observed an increased proportion of FLS2-FLS2 association from two but not more FLS2 molecules, which also exhibited longer-lasting association time, suggesting stochastic binding in seconds into dimer upon ligand elicitation. Our synthetically engineered receptor, designed to mimic natural dimerization, outperforms the unmodified monomeric FLS2 consistently in each step of immune signaling transduction. This enhancement likely facilitates the rapid formation of functional FLS2-BAK1 heteromer complexes, which is crucial for effective signaling. While natural FLS2-BAK1 complex formation is known to be robust and swift, our synthetic approach enforces this interaction through oligomerization, potentially accelerating signaling processes that would otherwise evolve more slowly through natural plant-microbe co-evolution.

Our engineered tetrameric FLS2 receptors exhibit reduced lateral mobility on the PM and significantly impaired endocytic internalization. This suggests that an optimal level of receptor oligomerization is crucial for balancing robust complex formation and the maintenance of receptor homeostasis in both resting and activated states. Although tetrameric FLS2 enhances specific PTI responses compared to monomeric FLS2 due to BAK1 clustering, it does not achieve the optimal activity as observed on dimeric FLS2. This reduced efficacy is likely because the constrained conformational flexibility of tetramer or higher oligomer destabilizes the assembly of multicomponent complexes. Additionally, the tetramer exhibits impaired endocytic internalization and degradation post-activation, possibly due to compromised compatibility with endocytic adaptors (*30, 40, 41*). This impairs the replenishment of newly synthesized FLS2 molecules and thus leads to defects in long-lasting innate immunity with multiple rounds of immune responses. In addition, our synthetic engineering has identified a dimeric resting-state FLS2 receptor that enhances defense responses against bacterial infections but does not affect plant growth at its resting state. This offers an effective strategy for improving plant defense capabilities.

Mammalian TCR dimerization is a common feature of TCR activation. As TCRs do not have intrinsic kinase activity, dimerization is indispensable to induce conformation change and, therefore, enable the recruitment of adaptor proteins and signaling kinases that form a signal hub to activate the immune signaling (*35, 42*). Unlike the mammalian TCR, plant PRRs possess kinase domains or can utilize the kinase domains of the constitutively associated LRR-RKs to directly mediate the trans-phosphorylation with their co-receptors to initiate immune signaling upon recognizing pathogenic signal (*3, 8, 9*). Therefore, a conformation change in the kinase domains of plant RKs and their co-receptors is not necessarily required as long as ligand binding brings those kinase domains into close proximity (*43*). Here, we found dimeric resting-state FLS2 still binds to its co-receptor BAK1 with the same affinity as monomeric-FLS2 and, therefore, clusters BAK1, which results in higher phosphorylation in each step of signaling transduction, leading to higher ROS production. Higher orders of FLS2 clusters, starting from the tetrameric assembly, showed a reduced probability of binding with BAK1 upon ligand perception but still induced BAK1 clustering and displayed enhanced immune activation. These results highlight potential mechanisms by which receptor clustering amplifies PTI signaling by enhancing the kinase activities of the receptor and co-receptor complex, thereby facilitating the phosphorylation of downstream signaling molecules. This is further supported by the observation that BAK1 clustering phenocopied FLS2 clustering in flg22-triggered ROS production. Indeed, the formation of high-order assembly is also a common strategy adopted by metazoan immune receptors to amplify the immune signaling, such as TCR activation nucleates the assembly of the Myddosome (*36, 42*), EGFR multimerization to cooperatively activate its kinase domain (*44*) and TLR3 oligomerization upon binding viral dsRNAs to enable cooperative interactions between the ligand and receptor (*45*). Receptor clustering locally concentrates the receptors and their associated signaling molecules, thereby providing a synergistic effect to increase the signaling sensitivity and strength, enhance signaling transduction, and also enable a spatial organization of these molecules (*46, 47*). The molecular activities of concentrated molecules are improved due to, for instance, the increased substrate concentration and molecular organization inside clusters (*11*), the high densities of molecular binding sites (*10*), and the enhanced binding kinetics and recruitment of downstream molecules (*48*), which thereby promotes the related signal propagation. The signal transduction in the FLS2/BAK1 complex and BIK1 is intricate and contains entangled multiple steps or mixed steps of auto- and trans-phosphorylation (*26, 27*). The dynamic interplays and signaling propagation between the clustered FLS2/BAK1 complexes and BIK1 upon ligand elicitation would need further study to be well-characterized.

## Acknowledgments

We are grateful to Prof. Dahai Luo (Nanyang Technological University, Singapore) for critical reading of the manuscript. We thank Yifan Cao for assistance in molecular clone and Rebecca Liew Hui Ting for helping on screening of transgenic plants. We thank Prof. Eui-Hwan Chung (Korea University, Korea) for sharing the *Pst* DC3000 strain and Prof. Xiufang Xin (Chinese Academy of Sciences, China) for sharing the *Pst* DC3000 D36E strain.

## Funding

Singapore Ministry of Education Tier 3 grant (MOE2019-T3-1-012), Research Centre of Excellence award to the Institute for Digital Molecular Analytics and Science (EDUN C-33-18-279-V12) and National Research Foundation Singapore (MOH-000955, NRF-NRFI08-2022-0012) to YM.

## Author contributions

Conceptualization: ZM, YM; Methodology: ZM, YX, CPC, CJH, YM; Investigation: ZM, YX, CPC; Visualization: ZM, YX, CPC; Funding acquisition: YM; Project administration: YM; Supervision: YM; Writing – original draft: ZM, YM; Writing – review & editing: ZM, YM.

## Competing interests

The authors declare that they have no competing interests related to this work.

## Data and materials availability

All data are available in the main text or the supplementary materials. Constructs and transgenic *Arabidopsis* lines are available on request.

## Notes

### Competing Interest Statement

The authors have declared no competing interest.

