## supplementary files for "Synthetic FLS2 receptor oligomer boosts plant innate immunity"

**The PDF file includes:**

Materials and Methods
Figs. S1 to S4

**Other Supplementary Materials for this manuscript include the following:**

Movies S1 to S3

### Materials and Methods

#### Plant growth

All the *Arabidopsis thaliana* lines reported in this study were in the Columbia ecotype (Col-0) background. The *fls2* (SALK_026801C) was obtained from the Arabidopsis Biological Resource Center (ABRC). To image the FLS2-CC-mG signal, the seeds were first surface-sterilized in 15% bleach (v/v) for 1 minute and washed with autoclaved water. Then, the seeds were plotted on half strength of Murashige and Skoog (MS) medium before growing in the growth chambers under long-day cycles (16 h light/8 h dark) at 22 °C. Five-day-old seedlings were used for elicitation and imaging. Two-week-old seedlings were used for WB assay to detect the MAPK activation and FLS2-BAK1 interaction. To test the immune activation-inhibited *Arabidopsis* growth, the indicated transgenic *Arabidopsis* seeds were surface-sterilized and plotted in the half strength of MS medium supplied with or without 100 nM flg22 and then further grew vertically under long-day cycles at 22 °C for five days before root length measurement. For ROS and bacterial infection assays, the seeds were directed grown in soil substrates (Peatmoss: Vermiculite: Perlite = 2: 1: 1) for 4-5 weeks under short-day cycles (10 h light/14 h dark) at 22 °C before leaf disc collection or bacteria injection. Transient expression was done in *N. benthamian*, grown in the same soil substrates for four weeks under long-day cycles at 24 °C before agrobacteria injection.

#### Plasmid construction

The binary vectors of multimeric FLS2 were generated through a customized gateway system. The DNA fragments of the FLS2 promotor and FLS2 gDNA sequence infused with 3xMyc tags were obtained from early reported plasmid *pFLS2::FLS2-3xMyc-GFP (33)* and respectively cloned to the gateway entry vector containing monomeric super-fold GFP (mG) to generate the gene expression set *pFLS2::FLS2-3xMyc-mG*. DNA fragments encoding dimer or tetramer CC motifs were then synthesized and inserted to construct *pFLS2::FLS2-3xMyc-Dimer-mG* and *pFLS2::FLS2-3xMyc-Tetramer-mG*, respectively. All the clones were done through homologous recombination by Exnase II (Vazyme). After sequencing verification, those gene expression sets were further cloned to the gateway destination vector pHGW to generate the binary vectors. Similar strategies were also applied to constructing the vectors of *pBIK1::BIK1-CC-mGs*, *35S::BAK1ECD-CC-BAK1KDs*, *35S::BAK1-mRuby2* and *35S::PEPR1-CC-mGs*. Here, the coding cDNA sequences of BIK1 and BAK1 were cloned from the *Arabidopsis* cDNA library and reported *pPZP212-pBAK1::BAK1-GFP* (*49*), respectively. *PEPR1* was cloned from the *Arabidopsis* cDNA library. For recombinant protein expression, the DNA fragments encoding CC motifs (from monomer to pentamer) and mG, separated by multiple linker residues, were integrated into the pET28a backbone through homologous recombination for bacteria expression. To generate the calibration standards *in vivo*, the above multiple *mG-CC* fragments were further fused with DNA fragments encoding membrane association peptide containing myristylation and palmitoylation (MAP) sites at N-terminus and then were integrated into binary vectors for plant expression. All the above binary vector plasmids were transformed into *Agrobacterium tumefaciens* strain GV3101 for either transient expression or *Arabidopsis* transformation through floral dip methods. Transgenic seeds with FLS2-CC-mG expression were screened by Hygromycin.

#### Transient expression

The *Agrobacterium* strains containing indicated binary vectors were cultured overnight at 28°C in LB broth medium supplemented with respective antibiotics. Afterward, the bacteria were collected and resuspended in the infiltration buffer (10 mM MES, pH 5.7, 10 mM MgCl2, and 100 mM acetosyringone) and incubated at 28°C for 2 h before being injected into the abaxial surface of *N. benthamian* leaves. The single-particle image was done at 16-20 h post *Agrobacterium* injection. For ROS and WB assays comparing multimeric engineering targets (FLS2, PEPR1, BAK1, BIK1), the initial *Agrobacterium* concentrations for injection were optimized to enable a comparable protein expression across different engineered proteins, which were further confirmed by WB. ROS and WB assays were done in the samples at 40-45 h post *Agrobacterium* injection.

**Bacterial inoculation**

Bacteria inoculation in four-week-old *Arabidopsis* seedlings was applied in this study to understand the bacteria invasion in multimeric FLS2 lines. For inoculation, $1X10^6$ CFU/mL of *Pseudomonas syringae* pv. tomato (*Pst*), DC3000 WT or $2X10^6$ CFU/mL of *Pst* DC3000 D36E strains, suspended in 10 mM $MgCl_2$, were injected into the abaxial surface of rosette leaves. The seedlings were then grown under normal growth conditions for another three days. The leaf tissues with bacteria inoculation were collected and ground in a bead beater to quantify the internal bacteria colonization. The tissue lysis was resuspended in 10 mM $MgCl_2$. With a series dilution, the tissue suspension was further dropped on NYG (3 g/L yeast extract, 5 g/L peptone, 20 g/L glycerol) agar (1.5%, W/V) plates which were further incubated at 28 °C chamber for another two days before the bacteria clone numbers were then counted.

**ROS assay**

The plant leaves from five-week-old *Arabidopsis* or *N. benthamian* with indicated protein transient expression were punched to get leaf discs of identical size (Diameter = 0.4 cm). The leaf discs were floated on sterile water in a 96-well plate (Greiner Bio-One GmbH) overnight under continuous light. To elicit ROS burst, the water was replaced by ROS elicitation solution (20 nM flg22/elf26/pep1, 20 mM luminal L-021, and 20 mg/mL horseradish peroxidase), and the luminescence was immediately monitored by a BioTek cytation 5 multimode reader (Agilent) with a time interval as 80 s. For each sample, the ROS production was recorded as relative light units (RLUs) from at least 6 leaf discs, respectively.

**Western blot assay**

Total proteins were extracted from *Arabidopsis* or *N. Benthamian* leaves with indicated protein transient expression. The collected tissues were frozen in liquid nitrogen and ground in a bead beater. The tissue powder was then resuspended in lysis buffer: 50 mM HEPES, pH7.4, 150 mM KCl, 1 mM EDTA, 0.5% Triton X100, 1 mM DTT, and pierce proteinase inhibitor (Thermo Scientific, 1 tablet for 50 mL solution). After removing the cell debris by centrifugation at 12,000 x *g* for 10 min at 4 °C, the supernatant was collected for western blot analysis. To analyze the FLS2 protein level in each multimeric FLS2 transgenic *Arabidopsis* line, FLS2 was detected by anti-Myc (9E10, 1:1000 dilution). To check the MAPK cascade activation, 2-week-old *Arabidopsis* seedlings grown in 1/2 MS medium were sprayed with 20 nM flg22 supplied with 0.01% (v/v) Silwet L-77. The samples were then collected at indicated time points post flg22 spray.

For PEPR1 transient expression in *N. benthamian*, the elicitor 20 nM pep1 was directly injected into the leaf, and the samples were then collected. Total proteins were further isolated using the same methods mentioned above. Phospho-p44/42 MAPK (Erk1/2) (Thr202/Tyr204) monoclonal antibody (Cell Signaling, 1:3000 dilution) was applied to detect the phosphorylated MAP kinases. To check the BIK1 phosphorylation, the membrane proteins were enriched by lipid microsome isolation. After removing the cell debris, the supernatant was further subjected to ultracentrifugation at 100,000 x *g* for 30 min at 4 °C to spin down the lipid microsomes, which were resuspended in lysis buffer supplied with 3% SDS. anti-BIK1 (Agrisera # AS164030) was used to detect the BIK1 level. The isolated proteins were separated in SDS-PAGE gel and transferred to the PVDF membrane (0.45 um) through wet transferring. The membranes were then blocked by 5% (w/v) BSA (Bio Basic) in TBST at room temperature for 1 h and further incubated with antibodies diluted in TBST buffer as mentioned above overnight in the cold room (4 °C). Afterward, the corresponding secondary antibodies conjugated with HRP were added and incubated with membrane at room temperature for 1 h. To detect the protein signal, chemiluminescence was induced by adding a luminol peroxide detection reagent (Amersham) and captured through the ChemiDoc image system (Bio-Rad). All western blot analyses were repeated at least two times, and similar results were obtained.

**Immunoprecipitation**

To analyze the FLS2 and BAK1 interaction, rosette leaves (~ 1 g) of five-week-old *Arabidopsis* seedlings were cut into lead stripes (width ~0.2 cm) and soaked in sterile water overnight under continuous light. 20 nM flg22 was then applied for 10 min before sample collection and total protein extraction. 1 mL of lysis buffer was added to each sample. After centrifugation at 12000 x *g* for 10 min at 4 °C, the supernatant (~1.2 mL) was incubated with 30 uL of GFP-Trap agarose beads (ChromoTek, #AB_2631357) for 1 h at 4 °C. The beads were further collected by centrifugation at 2500 x *g* for 5 min and washed three times with washing buffer (protein lysis buffer without adding detergent). In the last step, 100 mL of protein lysis buffer containing SDS loading dye was added to the bead pellet. Those proteins were boiled at 95 °C for 5 min to release the proteins and then subjected to gel separation and immunoblotting. FLS2 was detected by anti-Myc, and BAK1 was detected by anti-BAK1 polyclonal antibodies (Agrisera #AS121858, 1:3000 dilution). To detect the RbohD phosphorylation, the Flag-RbohD was enriched through immunoprecipitation by anti-Flag magnetic agarose (Thermo Fisher, A36798). Total and phosphorylated-RbohD were further evaluated by anti-Flag M2 antibody (Merck, #F3165) and anti-phospho-RbohD (S343, S347), respectively.

**Native PAGE gel electrophoresis**

Native PAGE gel was prepared according to the lab-optimized receipt. The stacking gel contains 0.35 M Tris-HCl pH 8.8, 4% (w/v) Acrylamide, 0.05% (w/v) ammonium persulfate, and 0.1 (v/v) % TEMED. The separation gel was prepared using the same receipt except for adding 9% (w/v) acrylamide. For gel electrophoresis, 4 ug of recombinant mG-CC proteins or 10 ug of total proteins extracted from *N. benthamiana* leaf cells with MAP-mG-CC transient expression were loaded and separated in the running buffer containing 250 mM Tris pH 8.3 and 192 mM glycine at 4 $^{o}$C with constant current 0.02 A. The mG signal was captured under UV.

**Fluorescent image and analysis**

To analyze the oligomeric status of FLS2/BAK1/PEPR1-CC-mGs. The FLS2/BAK1/PEPR1-CC-mGs and calibration standards from monomer to pentamer) were transiently expressed in *N. benthamian* leaves. At 16-18 h post *Agrobacterium* injection, the single particles of oligomerized mGs on the PM were recorded by VA-TIRFM integrated with a Nikon ECLIPSE Ti2-E inverted microscope (Nikon Instruments) equipped with a 100x (NA1.49) Plan-Apo objective lens (Nikon Instruments). VA-TIRFM was achieved through the iLAS 2 platform (Gataca systems). The mG signal was excited at 488 nm (Vortran), and the emission was collected at 505-545 nm by an ORCA-Fusion sCMOS camera (Hamamatsu). The excitation laser power and capturing exposure time were identical across different samples. All the image acquisitions were controlled by MetaMorph (Molecular Devices) software. To analyze the single-particle intensity of mG, the raw images were subjected to background subtraction in FIJI software, and the 'Threshold' function was then applied to make a binary image to highlight the mG single particle signal for the subsequent particle selection by using 'analyze particle' function. After adding all the particles into the ROI manager, the total intensity of each single particle was measured in FIJI. The single-particle intensities for each oligomerized FLS2/BAK1-CC-mG type were plotted and subjected to single- or multiple-peak Gaussian distribution fitting. The Peak values for calibration standards were then plotted to generate the calibration curve. The oligomeric status of FLS2/BAK1-CC-mGs was determined by comparing the peak values of FLS2/BAK1-CC-mG single particle intensities with the calibration curve.

The single particle dynamics in either *Arabidopsis* or *N. benthamian* leaf epidermal cells were recorded using the same instrument as above. Stream imaging was applied to capture the dynamic diffusion of FLS2/PEPR1-CC-mGs with an interval of 200 ms for at least 2 s. The single particle dynamics were quantified following our early report (*50*). Time-lapse imaging with an interval of 1 s for a total time range of 3 min was applied to record the FLS2 endocytic lifetime, which was further analyzed through the kymograph function in FIJI. For the dual-colour image of FLS2-CC-mGs and BAK1-mRuby2/FLS2-mCh in *N. benthamian* leaves. The mRuby2 or mCh (mCherry) signal was excited at 561 nm (Coherent), and the emission was collected at 589-625 nm. The emission of mRuby2/mCh and mG was split through a W-VIEW GEMINI-2C image splitting system, equipped with a dichroic beam splitter (Hamamatsu) dichroic mirror and directed into two detection arms terminating in two cameras, respectively. Time-lapse imaging with an interval of 1 s was applied to record the FLS2-FLS2 association with or without flg22 elicitation. The same *N. benthamian* samples were also used to measure the fluorescent lifetime of FLS2-mG. The FLIM image was done through the STELLARIS 8 FALCON (Leica) platform. The mG was excited at 488 nm with a pulse picker (pulse frequency 80 MHz), directed through a 63 x (NA1.2) water lens (Leica), and the emission photons were detected by Power HyD X detector (Leica). The fluorescent lifetime of FLS2-mG was monitored according to the emission photon decay.

To check the bulk FLS2/PEPR1-CC-mGs signal in *Arabidopsis* transgenic lines. Five-day-old *Arabidopsis* cotyledons were observed under a Nikon ECLIPSE Ti2-E inverted microscope (Nikon Instruments) equipped with CSU-W1 confocal spinning unit (Yokogawa) through a 100x (NA1.49) Plan-Apo objective lens (Nikon Instruments). The mG signal was excited at 488 nm (Vortran), and the emission was collected at 505-545 nm by an ORCA-Fusion sCMOS camera (Hamamatsu). To record the FLS2-CC-mG engaged endosomes at the indicated time points after flg22 elicitation, Z-stack images with 0.4 μm of step size were captured for a depth of at least 5 μm from the surface. Z-projection with maximum intensity was further applied to display the endosomes.

**Molecular simulation**

Computer simulation of the random 2D diffusion of protein particles interacting with prescribed inter-particle interactions was carried out using Large-scale Atomic/Molecular Massively Parallel Simulator (LAMMPS) molecular dynamics simulation code1(*51*). The simulation code uses dimensionless units σ for length and τ for time. The domain is 100σ × 100σ, where σ is defined to be 0.1 μm so as to represent a domain of 10 μm × 10 μm. Periodic boundaries are applied along each dimension. Two types of particles are randomly placed on the domain at a density of 70 particles in 100 μm2 for green particles and a density of 100 in 100 μm2 for red particles. The interaction energy between particles is van der Waals in nature, described by the Lennard-Jones potential with two parameters: the depth of the attractive energy well, ε, in kT units (k is Boltzmann constant and T is temperature in Kelvin) and a particle diameter taken to be 0.22 σ or 22 nm (10x smaller than the diffraction limit of visible light). Particle pairs of the same type (red-red or green-green) interact weakly with an attractive energy depth of 0.1 kT. On the other hand, the inter-particle attraction between particle pairs of different types (red-green) varies from 0.1 kT to 6 kT to simulate protein-protein association via binding. Simulations were each carried out for 1 million steps at a temporal step size of 0.005τ, and the trajectory data from the last 100,000 steps was used for analyses. An in-house MATLAB script was used to calculate the percentage of inter-particle association as $2n_{pair}/(n_{red} + n_{green})$, where $n_{pair}$ is the number of particle pairs within cutoff of 100 nm, and $n_{red}$ and $n_{green}$ are the numbers of red and green particles respectively.

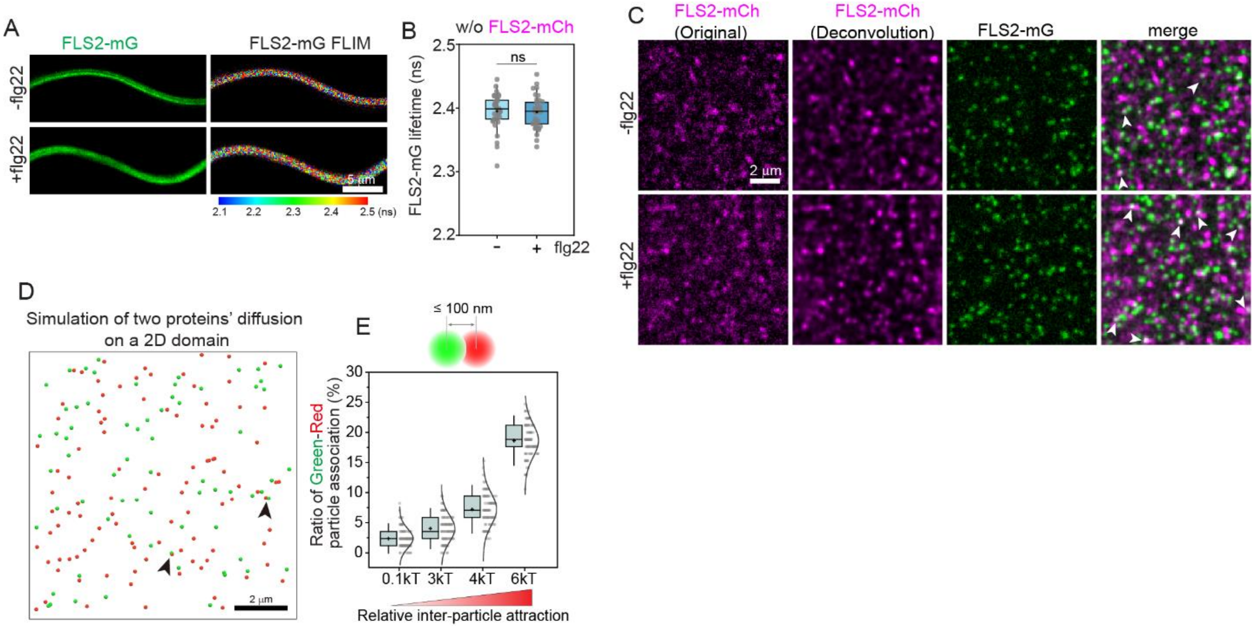


**Fig. S1. Single-particle and FLIM image for FLS2-mG/mCh.** (**A**) FLIM image for FLS2-mG in the absence of FLS2-mCh expression. Intensity and lifetime image of FLS2-mG with or without an elicitation by 20 nM flg22 for 5 min were shown. Colour bar indicates the corresponding lifetime. Scale bar, 5 μm. (**B**) Quantification of FLS2-mG lifetime in (A), n = 42 and 43 of ROIs from left to right. (**C**) Original images for shown in Fig. 1D. Noted the channel of FLS2-mCh was processed through a deconvolution algorithm for denoising the image. Scale bar, 2 μm. (**D**) Computer simulation to recapitulate two particles' overlapping on a 2D panel. The green and red particles with the same densities as FLS2-mG and FLS2-mCh single particles in Fig. 1D, respectively, were input into the simulation. Black arrowheads mark the green-red particle association. Scale bar, 2 μm. (**E**) Quantification of the ratios of green-red particle association as a function of relative inter-particle attraction as expressed in terms of thermal energy. Two particles with a distance lower than or equal to 100 nm were considered as an association. n=100 replicates. Scale bar, 2 μm. Noted 0.1 kT means the energy attraction is just 10% of the thermal energy, implying that thermal fluctuations dominate the particle. In this condition, inter-particle attraction is negligible and overlapping between particles occurs only incidentally. Box plots in (B) and (E) indicate mean (cross symbols), median (center bars), quartiles (box limits), and SD (whiskers).

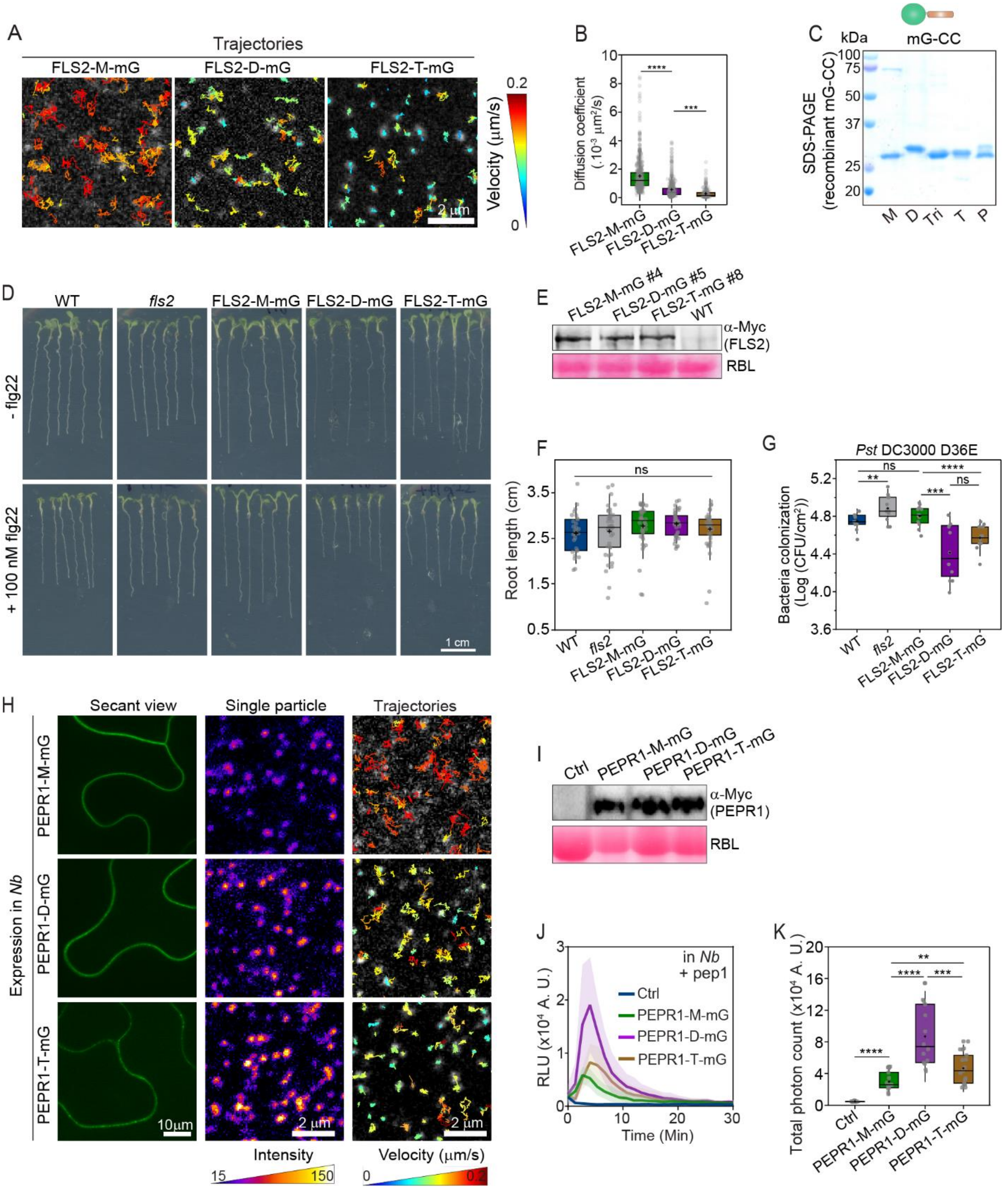
A
Trajectories
FLS2-M-mG
FLS2-D-mG
FLS2-T-mG
2 μm
Velocity (μm/s)
0.2
0
B
Diffusion coefficient (.10-3 μm²/s)
FLS2-M-mG
FLS2-D-mG
FLS2-T-mG
C
kDa
mG-CC
SDS-PAGE (recombinant mG-CC)
M D Tri T P
D
WT
fls2
FLS2-M-mG
FLS2-D-mG
FLS2-T-mG
- flg22
+ 100 nM flg22
1 cm
E
FLS2-M-mG #4
FLS2-D-mG #5
FLS2-T-mG #8
WT
α-Myc (FLS2)
RBL
F
Root length (cm)
ns
G
Pst DC3000 D36E
Bacteria colonization (Log (CFU/cm²))
H
Secant view
Single particle
Trajectories
Expression in Nb
PEPR1-M-mG
PEPR1-D-mG
PEPR1-T-mG
10μm
2 μm
Intensity
Velocity (μm/s)
I
Ctrl
PEPR1-M-mG
PEPR1-D-mG
PEPR1-T-mG
α-Myc (PEPR1)
RBL
J
in Nb + pep1
Ctrl
PEPR1-M-mG
PEPR1-D-mG
PEPR1-T-mG
RLU (x10⁴ A. U.)
Time (Min)
K
Total photon count (x10⁴ A. U.)
Ctrl
PEPR1-M-mG
PEPR1-D-mG
PEPR1-T-mG

**Fig. S2. Receptor clustering promotes plant PTI responses.** (**A**) Representative single-particle diffusion trajectories of FLS2-M/D/T-mG. Colour bar indicates the mean velocity of the trajectories. Scale bar, 2 μm. (**B**) Quantification of the single particle diffusion coefficient of FLS2-M/D/T-mG, n=561, 515 and 227 from left to right. (**C**) Coomassie blue-stained SDS-PAGE gel image of recombinant mG-CC proteins, following the CC valencies of Dimer (D), Trimer (Tri), Tetramer (T) or Pentamer (P). mG without CC domain was designated as Monomer (M). (**D**) Growth assay of WT, *fls2,* and FLS2-M/D/T-mG transgenic *Arabidopsis* lines in 1/2 MS medium supplemented with or without 100 nM flg22 for 5 days. Scale bar, 1 cm. (**E**) Immunoblot analysis by anti-Myc antibody to test FLS2 expression across selected FLS2-M/D/T-mG transgenic *Arabidopsis* lines. Ponceau S staining of rubisco large subunit (RBL) was used as a loading control. (**F**) Quantification of the root length in (E) with no flg22 addition. n= 29,42,42,42 and 38 of seedlings from left to right. (**G**) Quantification of internal bacteria colonization at 3 days post-infection. $2 \times 10^6$ CFU·mL$^{-1}$ of *Pst* DC 3000 D36E were injected into the rosette leaves of 4-weeks-old WT, *fls2* and FLS2-M/D/T-mG *Arabidopsis* seedlings for infection. n=12 replicates for each genotype. Error bars = SD. (**H**) Engineering and imaging the *Arabidopsis* PEPR1 receptor clusters in *N. benthamiana* leaves. The PEPR1 was fused with CC domains followed by a mG at its C-terminal like the same strategies applied on FLS2 (Fig. 2A). The single-particle images were recorded under VA-TIRFM, and their trajectories were extracted. Colour bars indicate the single particle intensity and diffusion mean velocity, respectively. Scale bar, 2 μm. (**I**) Immunoblot analysis to assess the PEPR1 expression in *N. benthamiana* leaves. The initial concentrations of *Agrobacterium* for injection into *N. benthamiana* leaves were adjusted to ensure a comparable expression across PEPR1-M/D/T-mG transient expression before the following ROS assay. PEPR1-M/D/T-mG were detected by anti-Myc antibody. Control (Ctrl) indicates the transient expression of mG alone. Ponceau S staining of RBL was used as a loading control. (**J**) Measurement of ROS in leaf discs of *N. benthamiana* with PEPR1-M/D/T-mG and Ctrl transient expression as in (I) and elicited by 20 nM pep1. Relative luminescence units (RLU) were monitored in real-time. Solid lines and shaded area represent mean ± SD from multiple leaf discs. (**K**) Quantification of total photon reading in 1-16 min in (L) representing the total ROS generation. n = 16, 15, 14 and 17 of leaf discs from left to right. Box plots in (B), (F), (G) and (K) indicate mean (cross symbols), median (center bars), quartiles (box limits), and SD (whiskers). Significant differences were determined via one-way ANOVA with multiple comparisons (****$p \leq 0.0001$, ***$p \leq 0.001$, **$p \leq 0.01$, ns = not significant).

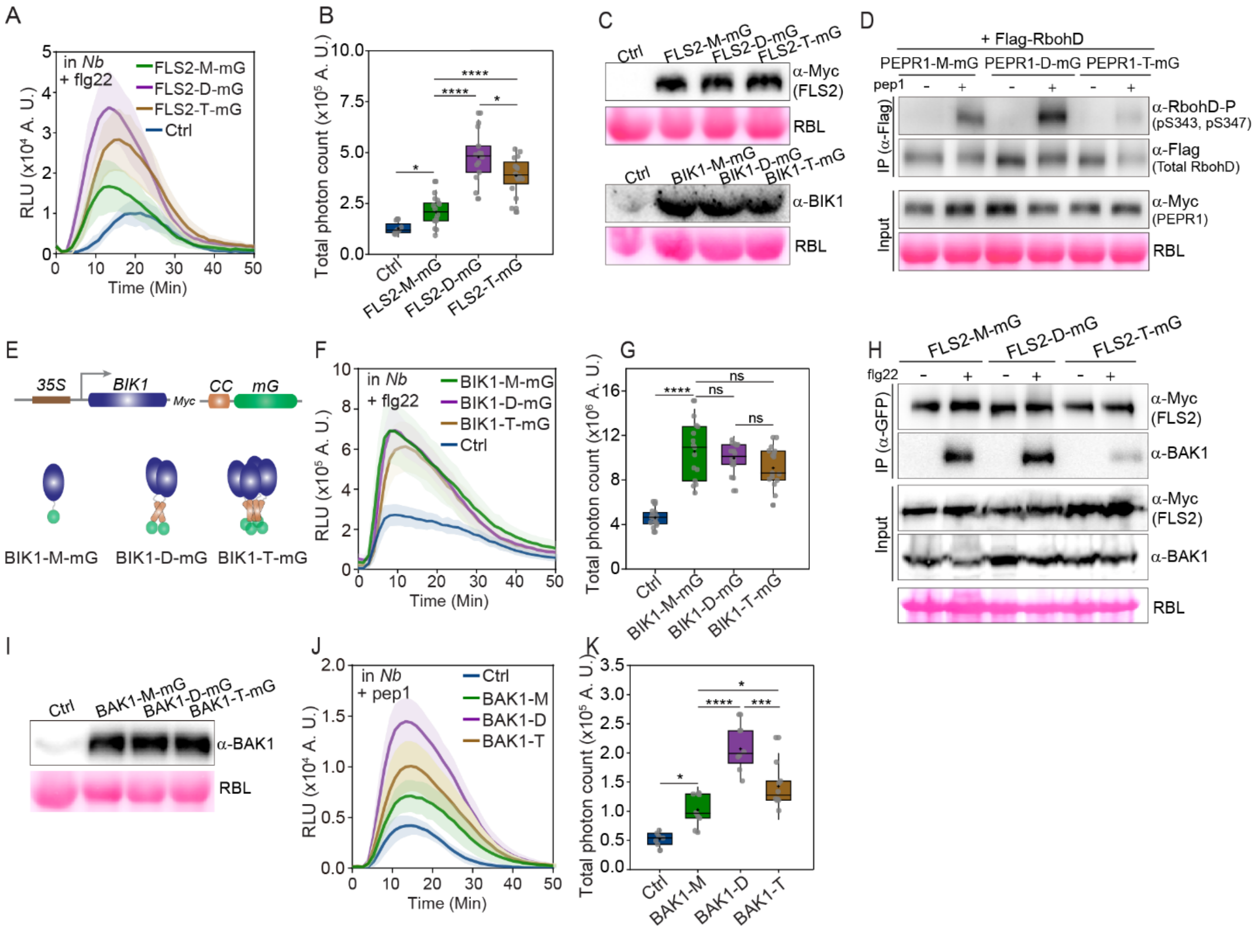


**Fig. S3. Dissect the PTI signaling pathway with engineered receptor clustering.** (**A**) Measurement of ROS in leaf discs of *N. benthamiana* with FLS2-M/D/T-mG or mG (Ctrl) transient expression and elicited by 20 nM flg22. RLUs were monitored in real time. Solid lines and shaded area represent mean ± SD from multiple leaf discs. (**B**) Quantification of total photon reading in 5-30 min in (A) representing the total ROS generation. n = 8, 14, 14 and 16 of leaf discs from left to right. (**C**) Immunoblot analysis to assess the FLS2 (detected by anti-Myc) and BIK1 (detected by anti-BIK1) expression in *N. benthamiana* leaves. The initial concentrations of *Agrobacterium* for injection into *N. benthamiana* leaves were adjusted to ensure a comparable expression across FLS2-M/D/T-mG or BIK1-M/D/T-mG transient expression before the following ROS assay. (**D**) RbohD phosphorylation in the presence of PEPR1-M/D/T-mG that were transiently co-expressed with RbohD-flag in *N. benthamiana* leaves and further elicited with or without 20 nM flg22 for 10 min. RbohD was enriched by anti-flag beads and then assessed through antibodies specifically detecting the phosphorylation (p) at serine 343 (S343) and S347. Ponceau S staining of RBL was used as a loading control. (**E**) Schematic illustration of BIK1 engineering for BIK1 clustering. 35S promoter-driven BIK1 was fused with CC domains, assembling to either Dimer (D) or Tetramer (T), followed by a mG at its C-terminal. Construct with no CC domain was designated as Monomer (M). (**F**) Measurement of ROS in leaf discs of *N. benthamiana* with BIK1-M/D/T-mG or mG alone (Ctrl) transient expression and elicited by 20 nM flg22. RLUs were monitored in real-time. Solid lines and shaded area represent mean ± SD from multiple leaf discs. (**G**) Quantification of total photon reading in 5-30 min in (E) representing the total ROS generation.

n = 15, 15, 16 and 15 of leaf discs from left to right. (**H**) FLS2-BAK1 interaction assessed by Co-IP assay. Leaf strips from 5-week-old *Arabidopsis* seedlings of FLS2-M/D/T-mG were elicited with or without 100 nM flg22 for 10 min. Immunoprecipitation was done by anti-GFP agarose beads. FLS2 and BAK1 were detected by anti-Myc and anti-BAK1 antibodies, respectively. Ponceau S staining of RBL was used as a loading control. (**I**) Immunoblot analysis to assess the BAK1 (detected by anti-BAK1) expression in *N. benthamiana* leaves. The initial concentrations of *Agrobacterium* for injection into *N. benthamiana* leaves were adjusted to ensure a comparable expression across BAK1-M/D/T before the following ROS assay. The expression of the empty vector was set as Control (Ctrl). (**J**) Measurement of ROS in leaf discs of *N. benthamiana* with BAK1-M/D/T or empty vector (Ctrl) transiently co-expressing with PEPR1-M-mG and elicited by 20 nM of pep1. Relative luminescence units (RLUs) were monitored in real-time. Solid lines and shaded area represent mean ± SD from multiple leaf discs. (**K**) Quantification of total photon reading in 5-30 min in (J) representing the total ROS generation. n = 7, 10, 8 and 10 of leaf discs from left to right. Box plots in (B), (G), and (I) indicate mean (cross symbols), median (center bars), quartiles (box limits), and SD (whiskers). Significant differences were determined via one-way ANOVA with multiple comparisons (****$p \leq 0.0001$, ***$p \leq 0.001$, *$p \leq 0.05$, ns = not significant).

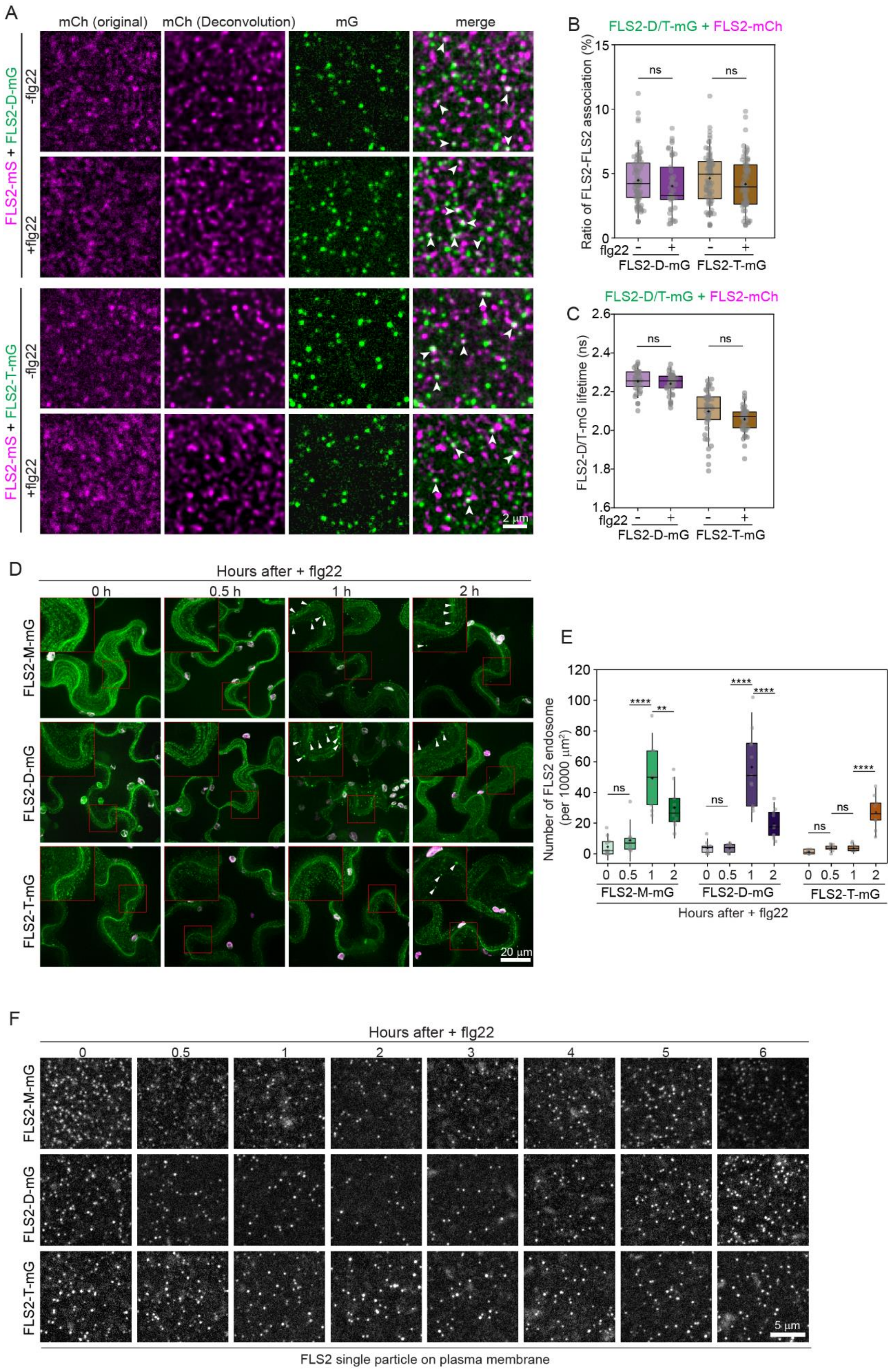
A
mCh (original)
mCh (Deconvolution)
mG
merge
FLS2-mS + FLS2-D-mG
-flg22
+flg22
FLS2-mS + FLS2-T-mG
2 μm
B
FLS2-D/T-mG + FLS2-mCh
Ratio of FLS2-FLS2 association (%)
ns
flg22
FLS2-D-mG
FLS2-T-mG
C
FLS2-D/T-mG + FLS2-mCh
FLS2-D/T-mG lifetime (ns)
D
Hours after + flg22
0 h
0.5 h
1 h
2 h
FLS2-M-mG
FLS2-D-mG
FLS2-T-mG
20 μm
E
Number of FLS2 endosome (per 10000 μm²)
Hours after + flg22
F
Hours after + flg22
5 μm
FLS2 single particle on plasma membrane

**Fig. S4. Image the dynamic turnover of FLS2 after elicitation.** (**A**) Single particle images of FLS2-D/T-mG co-expressing with FLS2-mCh in *N. benthamiana* leaves with or without an elicitation by 20 nM flg22 in 5 min. FLS2-mCh image was processed through a deconvolution algorithm for denoising. White arrowheads in overlapped images mark FLS2-D/T-mG associating with FLS2-mCh single particles. Scale bar, 2 μm. (**B**) Quantification of the ratio of single particle association in (A), the number of FLS2-D/T-mG and FLS2-mCh particles showing association was normalized by the total number of single particles from two channels. n = 37, 36, 36, and 36 of ROIs (region of 100 $\mu m^2$) from left to right. (**C**) Quantification of FLS2-D/T-mG lifetimes through FLIM image for FLS2-D/T-mG (as donor) in the presence of FLS2-mCh (as acceptor) in *N. benthamiana* leaves with or without an elicitation by 20 nM flg22 as in (**A**). n = 39, 40, 40 and 39 of ROIs from left to right. (**D**) Real-time image of FLS2 engaged endosomes in 5-day-old FLS2-M/D/T-mG *Arabidopsis* cotyledon epidermal cells after elicitation by 20 nM flg22. Projected images with a Z-stack of 5 μm in 0.4 μm of step size were shown. Zoomed images show 2 x enlarged areas in red boxes. Arrowheads mark the FLS2-engaged endosomes. Scale bar, 20 μm. (**E**) Quantification of the endosome numbers in each area of $1x10^4$ $\mu m^2$ in (C). n ≥ 7 ROIs for each panel. (**F**) A full-time-dependent map of FLS2 single particle numbers remaining on plasma membrane after 20 nM flg22 elicitation. Noted that the flg22 was washed out at 1 h post elicitation to avoid continuous elicitation. Scale bar, 5 μm. See also Fig. 4E for the quantification. Box plots in (B), (C) and (E) indicate mean (cross symbols), median (center bars), quartiles (box limits), and SD (whiskers). Significant differences were determined via one-way ANOVA with multiple comparisons (****p ≤ 0.0001, **p ≤ 0.01, ns = not significant).

**Movie S1. Real-time recording of FLS2-mG and FLS2-mCh on the PM at resting state.** FLS2-mG and FLS2-mCh were transiently expressed in *N. benthamiana* leaves and injected with water as a control before imaging under VA-TIRFM. Dual-channel simultaneous imaging was applied to record the dynamic diffusion of FLS2-mG and FLS2-mCh on the PM with a time interval of 1 second (Sec). White arrowheads mark two cases of FLS2-mG and FLS2-mCh association. Scale bar: 2 μm.

**Movie S2. Real-time recording of FLS2-mG and FLS2-mCh on the PM in the presence of flg22 elicitation.** FLS2-mG and FLS2-mCh were transiently expressed in *N. benthamiana* leaves and injected with 20 nM flg22 before image under VA-TIRFM within 5 min post injection. Dual-channel simultaneous image was applied to record the dynamic diffusion of FLS2-mG and FLS2-mCh on the PM with a time interval of 1 second (Sec). White arrowheads mark two cases of FLS2-mG and FLS2-mCh association. Scale bar: 2 μm.

**Movie S3. Tracking of FLS2-M/D/T-mG single-particle diffusion dynamics on the PM.** Stream image for 1 min with an exposure time of 500 ms was applied to record FLS2-M/D/T-mG single-particle diffusion under VA-TIRFM in the cotyledon epidermal cells of 5-day-old *Arabidopsis* seedlings. Single particle trajectories were extracted through the Trackmate plugin in FIJI and colored from red to blue indicating the relative highest to lowest diffusion velocity, respectively. Scale bar: 2 μm.
